# Probing behavior of *Diaphorina citri* (Hemiptera: Liviidae) on Valencia orange influenced by sex, color, and size

**DOI:** 10.1101/2019.12.24.885178

**Authors:** Timothy A Ebert, Michael E Rogers

**Affiliations:** University of Florida, Citrus Research and Education Center, Department of Entomology and Nematology, 700 Experiment Station Rd., Lake Alfred, FL 33850

**Keywords:** Electropenetrography, electrical penetration graph, EPG, negative results, Asian Citrus Psyllid

## Abstract

*Candidatus* Liberibacter asiaticus is vectored by the psyllid *Diaphorina citri* Kuwayama (Hemiptera: Liviidae) and putatively causes Huanglongbing disease in citrus. Huanglongbing has reduced yields by 68% relative to pre-disease yields in Florida. Disease management is partly through vector control. Understanding vector biology is essential in this endeavor. Our goal was to document differences in probing behavior linked to sex. Based on both a literature review and our results we conclude that there is either no effect of sex or that identifying such an effect requires a sample size at least four times larger than standard methodologies. Including both color and sex in statistical models did not improve model performance. Both sex and color are correlated with body size, and body size has not been considered in previous studies on sex in *D. citri*. An effect of body size was found wherein larger psyllids took longer to reach ingestion behaviors and larger individuals spent more time ingesting phloem, but these relationships explained little of the variability in these data. We suggest that the effects of sex can be ignored when running EPG experiments on healthy psyllids.

---

*Diaphorina citri* Kuwayama (Hemiptera: Liviidae) is the insect vector of *Candidatus* Liberibacter asiaticus (*C*Las) which is the putative causal agent of Huanglongbing disease. The disease and psyllid are now present in the three major citrus growing regions of the United States: Florida (Halbert 2005), Texas (Kunta et al. 2014), and California (Kumagai et al. 2013). The disease has reduced production in Florida from 291.8 million boxes in 2003-04 season down to 94.2 million boxes in the 2015-16 season (https://www.nass.usda.gov/Statistics_by_State/Florida) and threatens a similar outcome in the other regions. Worldwide, CLas is considered the second most important insect vectored bacterial plant pathogen, behind *Xyllela fastidiosa* (Mansfield et al. 2012).

One of the three foundations for managing *C*Las is through insecticide applications to reduce pest populations (Grafton-Cardwell et al. 2013). While mortality is an essential component of efficacy, what the insect does in the time between exposure and death can influence the overall success of the insecticide application as a disease management tool. Because *C*Las is phloem limited, management techniques for *D. citri* need to focus on the ability of the pesticide to reduce or prevent the psyllid from salivating into the phloem or ingesting phloem wherein inoculation and acquisition events occur respectively (Bonani et al. 2010, Serikawa et al. 2013). In studying this system, experiments that ignore sex risk missing important differences between the sexes but including sex as a planned treatment doubles the size of an experiment. The third option is to test one sex and assume that the other sex behaves differently in some unknown but significant way that will be investigated in future research. Understanding the behavioral differences in probing behavior between the sexes is essential for proper experimental design and biological interpretation of the results.

Many insects show differences between males and females beyond the required anatomical and genetic differences. Sex differences can influence vector competence as demonstrated in the following examples. *Bemisia tabaci* (Gennadius) (Hemiptera: Aleyrodidae) females of biotype Q were better than males at transmitting tomato yellow leaf curl virus, and the females were better than either males or females of biotype B. There was no significant difference between males and females of biotype B. (Ning et al. 2015). In fourteen populations of *Frankliniella occidentalis* Pergande (Thysanoptera Thripidae), males were more successful at transmitting tomato spotted wilt virus because females feed more intensively than the males, and females move less (van de Wetering et al. 1998). In *Macrosteles quadrilineatus* (Hemiptera Cicadellidae) females are better at transmitting the aster yellows phytoplasma in lettuce, and the pattern of disease spread for females was significantly more clustered (Beanland et al. 1999). In *Scaphoideus titanus* Ball (Hemiptera: Cicadellidae), males spend more time probing than females. Males and females took the same time to reach xylem, but males stayed in xylem longer. Females took longer to reach phloem than males, but males remained in phloem twice as long as females (Chuche et al. 2017). As the authors point out, this system could result in sex-related differences in acquisition or transmission. Such examples show that keeping track of males and females can be important in understanding the biology of the pest and its role as a vector. The question is whether or not it makes a difference in *D. citri.* A more general literature search was done to provide a background. Several reviews of *D. citri* biology already exist (Halbert and Manjunath 2004, Yang et al. 2006, Grafton-Cardwell et al. 2013, Hall et al. 2013, Stelinski 2019). While these reviews discuss some sex-related differences, sex was not their focus.

## Sex Ratio

The male-female ratio is often close to 50:50 (Tsai and Liu 2000, Nava et al. 2007, Hall et al. 2008, Hall and Hentz 2016, Hall 2018). However, sometimes there exists a greater percentage of females (Catling 1970, Pande 1971, Xu et al. 1994, Alves et al. 2014).

## Morphology

Females are generally larger than males. Female wings are longer (Martini et al. 2015). The host plant affects morphological variability in males and females, though the males tend to be more variable (Garcia-Perez et al. 2013).

The morphology of the antennae is similar between males and females, but there are short apical setae with tips that are recessed inward in the females but not the males (Onagbola et al. 2008). Females, but not males, have circumanal glands that produce aliphatic hydrocarbons, fatty and ester waxes that are excreted with the honeydew (Ammar el et al. 2013). Males produce a clear honeydew that is deposited on the leaf surface behind the insect. Females produce honeydew pellets that are propelled away from the insect and off the plant (Ammar el et al. 2013).

## Life History

Females live longer on average, though exact estimates depend on the host plant. On a curry leaf host, males live an average of 15 days versus 17 days for females, but on acid lime, the values are 17 and 20 respectively (Chakravarthi et al. 1998). On Rangpur lime, the mean values are 24 and 31 days while the difference is 21 and 31 days respectively (males vs females) on Sunki mandarin (Nava et al. 2007). Sometimes there is no significant difference with both surviving an average of about 30 days on a variety of hosts (Alves et al. 2014) up to 47 days on others. In starvation tests, males and females have equivalent survival rates when stressed by high temperatures and different relative humidity levels with LT_50_ values of 24.9 to 43.2 h at 25 C and 7 to 97% relative humidity respectively and 12.1 to 32.2 h at 30 C and 7 to 97% relative humidity respectively (McFarland and Hoy 2001). For comparison, the longest survival in starvation was 94.5 h and the longest survival in non-starvation is 117 days (Liu and Tsai 2000).

It takes the same number of days from egg to maturity for males and females (Nava et al. 2010, Hall and Hentz 2016). Development time is slightly shorter for males, but the proportion of females exceeds that of males at about 5 days after the first adults start to emerge from a cohort (Cifuentes-Arenas et al. 2018). The effect this has on psyllid biology may be mitigated by multiple overlapping generations in the field. Sexual maturity is reached by males and females 2-3 days post-eclosion (Wenninger and Hall 2007).

## Physiology

In uninfected adults, there was no difference between male and female levels of glutathione S-transferase activity. However, enzymatic activity was higher in females but not males when the adults were carrying *C*Las. In the same study, cytochrome P_450_ levels for males and females were roughly equivalent, but there was a significant interaction effect wherein the difference in cytochrome levels between infected and uninfected females was significantly different, but was only marginally significant for males (Tiwari et al. 2011). Acetylcholinesterase levels were the same between males and females, but the infection status of field-collected material was not determined (Tiwari et al. 2012).

There may be a male-female difference in response to some insecticides. Females but not males were repelled by pyrmetrozine deposits at 24 and 48 h (Boina et al. 2011).

## Immune Response

Males had higher mortality than females when infected with *Bacillus subtilis* via septic injury (Arp 2017). No difference due to sex was observed when bacteria were administered orally. Additional testing was done wherein *D. citri* were first primed using an artificial diet containing dead bacterial cells. These adults were then exposed to an artificial diet containing *Serratia marcescens*, and mortality was assessed until all insects died. Mortality rates were significantly lower for primed females relative to primed males (Arp et al. 2017).

## Dispersal

Males and females increased dispersal rates to the same extent as a function of increasing crowding from conspecifics (Martini et al. 2015). The flight distance and duration are the same for males and females (Arakawa and Miyamoto 2007, Martini et al. 2014a). At distances up to 60 meters, the numbers of dispersing males versus the numbers of dispersing females were the same at one, two, or three meters above the soil surface (Hall and Hentz 2011).

## Olfaction

Females avoid the odor of other females in a density-dependent manner, but odors from plants damaged by other psyllids were attractive and attraction increased with higher numbers of psyllids (Martini et al. 2014b). Thus highly damaged hosts are preferred, but within such a host the shoots with the fewest psyllids are selected (Martini et al. 2014b). Both males and females respond to female odors in a dose-dependent fashion, but neither respond to the odor of males (Martini et al. 2014b). Cuticular hydrocarbons are one source of odors, and males have hexadecane, dodecanoic acid, and tetradecanoic acid while these are below detectable levels in females. Females have acetic acid, ethyl undecanoate, hexanoic acid, geranyl acetone, decanoic acid, isopropyl tetradecanoate, 1-methylpropyl dodecanoate, and pentadecanoic acid that are below detectable levels in the male. Both males and females had 1-dodecanol. Males respond to female cuticular extract, but no other response was detected: female-female odor, male-male odor, or female-male odor (Mann et al. 2013). Mann et al. (2013) also field-tested the result using both clear and yellow sticky traps. More psyllids were captured in scented clear traps than unscented traps and a larger proportion of captured insects were male. For yellow traps, the total catch was the same for scented and unscented traps, but scented traps captured more males. A behavioral assay in the lab indicated that dodecanoic acid was the active compound. While a field assay with this compound captured more males at the highest use rates, the total catch was no different from the unscented control (Mann et al. 2013). The finding that males are attracted to females but the other interactions are not statistically significant has also been reported elsewhere (Gharaei et al. 2014).

Odors can be used to identify host plants. Both sexes are attracted to Mexican lime. Undamaged grapefruit attracted only females. Valencia and sour orange were not attractive to either sex. However, the general trend was for the female to find a suitable host, and the male is then attracted to the female (Gharaei et al. 2014). Females respond to plant volatile cues while males do not (Beloti et al. 2017). However, in one test it was the males that had a stronger preference for plant odors (Patti and Setamou 2010). The response to plant volatiles was affected by mating status, sex, and the presence of visual cues (Wenninger et al. 2009b). Unmated individuals are less responsive to plant volatiles in the absence of visual cues. The female antenna may be more sensitive to plant volatiles than the male antenna. The female was behaviorally responsive to plant odors in a Y-tube olfactometer test when the visual cue was a yellow sticky card and Navel, Sour orange, or *Murraya panniculata* L. Jack versus a clean air control, while there was no response in the male when given the same choice (Wenninger et al. 2009b). However, the response of individuals to specific odors is complex because the psyllid’s attraction to specific odors is partly a learned response (Stockton et al. 2017a, Stockton et al. 2017c, Stockton et al. 2017b).

Sex has a variety of effects on chemoreception and biosynthesis, most notably in roles related to pheromone production and detection. Several transcripts have been identified with putative roles in male detection of female pheromones (Wu et al. 2016b).

## Visual response

Fewer males responded to white or UV-blocked white light. However, within the responders, there was no difference in preference for one color over another for males versus females (Paris et al. 2017).

## Vibrational communication

*Diaphorina citri* uses vibrational communication by vibrating their wings to generate a signal that is transmitted through the tree. The frequency of the vibrations was negatively correlated with body size, but the relationship was statistically significant only for males. While both males and females signal, the male searches for the female who remains stationary (Wenninger et al. 2009a). In exploring the potential use of vibrational signals to trap male psyllids it was reported that females were more likely to respond to signals from other females relative to synthetic signals or white noise (male signals not tested), while males would respond to male or female signals and a synthetic signal equally and only ignore white noise (Rohde et al. 2013).

## Endosymbionts

There may be an interaction between temperature and sex of the psyllid in the abundance of endosymbionts. A Chinese colony of *D. citri* and *Candidatus* Carsonella ruddii showed no temperature effect in endosymbiont levels for females, but the abundance of the endosymbiont was reduced in males exposed to 5 °C relative to 27 and 40 °C. The colony from Pakistan showed no effect of temperature (Hussain et al. 2017). In both cases, males seemed to have lower levels of Carsonella, but the difference was not significant. In the same study, temperatures of 40 °C reduced levels of Wolbachia in females, for both the Chinese and Pakistani colonies. The same outcome was reported for males from Pakistan, but the lowest level of Wolbachia in the Chinese colony was at 5 °C (Hussain et al. 2017).

## *Candidatus* Liberibacter asiaticus

Exposure of nymphs to *Candidatus* Liberibacter asiaticus (*C*Las) infected plants increased male dispersal, but not female dispersal. Furthermore, it was more likely that dispersing males carried *C*Las (Martini et al. 2015). However, infection rates between males and females are the same (Martini et al. 2015, Wu et al. 2018a, Wu et al. 2018b). It has been reported that titers of *C*Las fluctuate with higher levels in autumn compared to spring for females, but the titers were not significantly different for males (Wu et al. 2018a). However, it has been observed that females have a higher transmission rate (26%) compared to males (18%) (Wu et al. 2016a).

*C*Las was present in the ovaries, but not testes, though in other respects *C*Las distribution was the same in males and females (Ammar et al. 2011a, Ammar et al. 2011b).

## Probing Behavior

Several manuscripts describe sex-related differences in probing behavior in *D. citri*. However, some only used females (Bonani et al. 2010, Zhu et al. 2010, Youn et al. 2011, Cen et al. 2012, Serikawa et al. 2012, Serikawa et al. 2013). Others either assumed that there was no difference or assumed a stable sex ratio and were interested in an average outcome (Boina et al. 2011, Yang et al. 2011, Kim 2013, Ebert and Rogers 2016). Of those manuscripts that looked for a sex difference in probing behavior, two manuscripts reported no significant differences between males and females (Luo et al. 2015, George et al. 2017). However, significant differences have also been found. The duration of phloem ingestion was significantly longer for females. The probe duration per event was shorter for females. The pathway duration per probe was longer in females. The number of phloem ingestion events was greater in females (Serikawa 2011). In another report, the minimum duration of phloem salivation was shorter in females than males, but in other respects, males and females were similar (Wu et al. 2016a).

## Color

Diaphorina citri has different color forms that include green, blue, red, orange, grey, and brown. Color differences were first documented over 90 years ago (Husain and Nath 1927). Abdominal color influences mating success and fecundity (Wenninger et al. 2009c, Stockton et al. 2017b). While blue males had the same size testes as orange males, females mated to orange males laid twice as many eggs relative to blue males. Experienced females avoid blue males (Stockton et al. 2017b). Body size is correlated with both sex and abdominal color with higher reproductive potential in blue/green individuals (Wenninger and Hall 2008, Wenninger et al. 2009c). It has also been shown that the blue/green morph was more capable of long-distance dispersal than the brown morph (Martini et al. 2014a). Finally, consistent with the observation that the blue/green morph had higher expression of CYP4 genes (Tiwari et al. 2013), insecticide applications increase the proportion of the blue/green morph (Martini et al. 2016).

Psyllids can change color over their lifetime based on age, mating status, sexual maturity, and possibly other factors. However, this change is not consistent enough to be diagnostic (Wenninger and Hall 2008). Gray/brown males and females are of roughly equal size, but they are smaller than any other sex/color combination. Gray/brown females were ca 30% smaller than blue/green females, while the difference between gray/brown versus blue/green males was ca 18% (Wenninger and Hall 2008). Both color and sex influenced the expression of the *Krüppel-homolog1* transcription factor involved in vitellogenesis and oogenesis where green females had higher relative expression levels than brown females and levels declined with female age from 1 to 14 days. There was no significant difference between green and brown males, but levels increased with age (Ibanez et al. 2019).

## Probing behavior (EPG)

This paragraph is an introduction to probing behavior in *D. citri* as measured using EPG. Probing behavior is a sequence of events resulting in nutriment acquisition or pathogen transfer. Specific events can be identified by repetitive patterns in the EPG output. These events are waveforms that are given abbreviations to aid in data entry, analysis, and presentation. The psyllid maintains osmotic balance by ingesting xylem (waveform G). Xylem ingestion also dilutes ingested toxins and aids in excreting toxins or their metabolites. The psyllid gets most of its nutrition from ingesting phloem (waveform E2). Once it has contacted phloem or xylem it has demonstrated that it is capable of further contact. Host plant acceptance is indicated by long phloem ingestion events (greater than 10 minutes). Spending a long time in xylem may indicate problems with osmotic balance or elevated levels of plant defensive compounds. More time spent until the first probe might indicate problems with host identification, while a long time to first phloem ingestion could indicate physical barriers like phloem fiber cells or might indicate difficulties recognizing the cues for locating the phloem or xylem. Once the psyllid has contacted the phloem or xylem it has demonstrated skill at reaching nutriment. However, if the nutriment is of poor quality the psyllid might try other feeding sites resulting in more probes after the first contact. Additionally, the insect is a vector, and transmission occurs during phloem salivation (E1), and maybe during phloem contact (waveform D). The acquisition of the pathogen by the insect is during phloem ingestion. As a final consideration, the response in the psyllid could be a shift in the mean, or it could be a change in the variability about the mean. These considerations are the conceptual beginning to converting six waveforms (non-probing NP, and five probing waveforms consisting of pathway C, xylem ingestion G, contact with phloem D, phloem salivation E1, and phloem ingestion E2) into a multitude of different variables (Tjallingii 1995, Backus et al. 2007, Sarria et al. 2009, Bonani et al. 2010, Ebert et al. 2015, Ebert et al. 2018b).

Given the inconsistent reports of a difference in the probing behavior between male and female *D. citri*, we tested for a difference in probing behavior due to sex. The issue is important in understanding the biology of this insect and its role as a vector. Likewise, it is important in designing other experiments where a difference between males and females could confound the results if sex differences are ignored. The review showed that sex, color, and size are significantly related to one another. Therefore, these factors were included in this study to identify which factor was more influential on the expected changes in feeding behavior. We also considered the possibility that the expected difference between males and females would be an artifact of associated differences in body size with females generally being larger than males.

## Materials and Methods

Psyllids in the colony were placed into two color groups blue+green versus grey+brown as done elsewhere (Wenninger and Hall 2008, Wenninger et al. 2009c, Wenninger et al. 2009b, Ibanez et al. 2019). This minimizes the subjective issue in placing individuals of mixed color into a discrete color group.

### Experimental design

The sample size for different conditions was male and green (MG=17), male and brown (MB=33), female and green (FG=20), and female and brown (FB=27). After recording, each psyllid was decapitated and the metathoracic leg removed. The head and leg were mounted on a slide using a coverslip and mounting media (Richard-Allan Scientific #4112, Kalamazoo, MI). Measurements were taken using a digital camera (OMAX 14MP A35140U, Omaxmicroscope.com) on a Wild M3Z microscope (www.leicabiosystems.com) and processed using ImageJ (version 1.51k at https://imagej.nih.gov/ij/). The head capsule was measured as the maximum distance from the outer margin of one compound eye to the outer margin of the other eye. The metathoracic femur length was also measured. There is a well-documented relationship between size, weight, and fitness in insects (Calvo and Molina 2005, Pekkala et al. 2011). Examining multiple measures of body size as well as their interaction better describes size. This is because many organisms undergo allometric growth wherein some body parts change faster than others. For a discussion of allometry see (West et al. 1997, Centorame et al. 2019, Patel and Matheson 2019, Sober et al. 2019). The simplest measure of size would have been dry weight, but we lacked the necessary equipment to accurately measure individual dry weights.

### Plants

We used *Citrus sinensis* (L. Osbeck.) Valencia orange scion on Kuharske citrange (*C. sinensis* X *Poncirus trifoliate* (L.) Raf.) rootstock (Southern Citrus Nurseries Inc., Dundee, FL: certified *C*Las free) for both rearing the psyllids and for the experiment. Plants were in 3.92 L black plastic pots filled with Fafard Professional Custom Mix (Agawam, MA). Plants were pruned to a height of 51 cm from the soil surface. Artificial light was provided by high-pressure sodium lamps (16:8 h L:D). Plants were fertilized (variously with Chelated Citrus Nutritional Spray from Southern Agricultural Insecticides Inc. Palmetto, FL, USA, Miracle-Gro All-Purpose Plant Food, Scotts Miracle-Gro, Marysville, OH, USA, or Harrell’s Profertilizer 12-3-8, Harrell’s LLC, Lakeland, FL) consistent with label instructions to promote flush.

### Insects

Psyllids were obtained from a *C*Las-negative colony maintained in the laboratory at the Citrus Research and Education Center, Lake Alfred, FL. The colony was tested periodically for *C*Las using PCR methods (Li et al. 2006) but has never tested positive (results not shown).

### Electropenetrography

All insects were recorded feeding on the abaxial surface of immature leaves (Ebert et al. 2018b). We used two 4-channel AC-DC monitors (EPG Technologies, Inc., Gainesville, FL, USA) in DC mode 150 mV substrate voltage. Data was acquired through a DI710 AD converter (Akron, OH) using Windaq software at a sampling rate of 100 Hz/channel. Psyllids were tethered using a 2 cm long 25.4 μm diameter gold wire (Sigmund Cohn Corp., Mt Vernon, NY) attached to thoracic tergites using silver glue (1:1:1 www, white glue:water:silver flake [8-10 μm, Inframat Advanced Materials, Manchester CT]). All head amps were set to an impedance of 10^9^ Ω. A 10 cm length of 2 mm diameter copper wire was inserted into the water-saturated soil as the positive electrode. There was a 30 m starvation period from the time the insects were removed from the colony until they were placed on the plant. All insects were wired during this period. Psyllids were not chilled or anesthetized with CO_2_. The recording was started before psyllids were placed on the plant to ensure that all recordings started in the NP behavior. Recordings were 23 h in duration. We chose to measure only the original six waveforms (NP, C, D, E1, E2, G) (Bonani et al. 2010). Lighting was provided by overhead fluorescent lights (24:0 h L:D). Room temperature was maintained at 26.6 °C. This was a standard protocol in this laboratory that has been published previously (Ebert and Rogers 2016, Killiny et al. 2017, Ebert et al. 2018b, Shugart et al. 2019).

### Data Analysis

The six recorded waveforms from each insect were used to calculate 89 variables that can be roughly described as durations, counts, percentages, the time to key events, and behaviors conditional upon some event (Ebert et al. 2015). If the null hypothesis is true (there are no treatment differences), then it is inevitable that some type I errors will be made. With the assumption that all the tests are independent, the probability of finding *r* Type I errors in *N* tries given a significance threshold of 0.05 is calculated as 0.95^(*N*−*r*)^ × 0.05^*r*^ × *N*!/(*r*! × (*N*−*r*)!). One should ask if it is reasonable to suppose that one or two significant outcomes in many tries (89 herein but varies by manuscript) represents a biological effect or a type I error. While considering the number of significant outcomes relative to the number of tests is one approach, a more standard solution to this problem is the Bonferroni correction wherein one divides the significance threshold by the number of tests (Rice 1989).

All variables discussed in this manuscript are calculated for each insect. We used the name of this variable even after statistical analysis. However, the value displayed no longer matches the name. Thus, the total duration of phloem salivation for each insect becomes a mean when these totals are averaged across insects. An elegant naming system was proposed to solve this issue for non-sequential variables (Backus et al. 2007), but the naming conventions become less user-friendly when applied to many of the variables used for aphids: see variable names in (Tjallingii 1995, Sarria et al. 2009, Tjallingii 2019).

Ebert 2.0 (http://www.crec.ifas.ufl.edu/extension/epg/sas.shtml) was used to analyze the data (Ebert et al. 2015, Ebert and Rogers 2016, Ebert et al. 2018b). All treatments are binary (male-female or green-other) therefore significance is tested by model F-test or parameter t-test as implemented in proc Glimmix in SAS (SASInstitute 1988). The use of mixed-model ANOVA results in improved power of statistical tests compared with non-parametric ANOVA (Gbur et al. 2012). In general, counts were square-root transformed, percentages were logit transformed, and everything else was log(variable + 0.1) transformed. These transformations were done with the sole purpose of improving the statistical models based on a graphical analysis of the residuals conforming to model assumptions of normality and homoscedasticity.

Estimates of sample size were done in Rstudio (version 1.1.456) running R (version 3.5.1) using the “pwr” package and the syntax pwr.t.test (d = (mean1-mean2)/pooled standard deviation, sig.level = 0.05, power = 0.80, type = ‘two.sample’). This sample size is an estimate of the required sample size assuming that the observed mean and standard deviation in the existing data are the correct values for the population.

The EPG recordings are published in DataVerse along with subsequent steps in the analysis. The programs are also included, though they only modifications from the originals involved identifying where to read and save files along with code specific to analyzing the data. The code for calculating the EPG variables is unchanged from previously published code (https://crec.ifas.ufl.edu/extension/epg/).

## Results

### Sex only

There were four variables that showed a significant difference between males and females: the percent of probing time spent in pathway (*df* 1, 86, *F*=4.21, *P*>*F* 0.0432), the standard deviation of the mean duration of xylem ingestion (*df* 1, 70, *F*=4.72, *P*>*F* 0.0333), the number of probes after the first xylem ingestion event (*df* 1, 86, *F*=4.01, *P*>*F* 0.0483), and the total duration of both phloem salivation and phloem ingestion for each insect (*df* 1, 64, *F*=4.4, *P*>*F* 0.0398). In 89 tries, there was a 0.46 probability of finding five or more significant outcomes. The p-value for all models was close to the 0.05 cut-off for significance, and no model was significant after applying a Bonferroni correction (0.05/89=0.000562).

### Sex and Color

There were three significant models with both sex and color. The mean duration of pathway (*df* 2, 85, *F*=5.48, *P*>*F* 0.0058, sex *P*>*F*=0.44, color *P*>*F*=0.002), The standard deviation of the mean duration of pathway (*df* 2, 85, *F*=3.72, *P*>*F* 0.0282, sex *P*>*F*=0.26, color *P*>*F*=0.019), and the number of probes after the first xylem ingestion event (*df* 2, 85, *F*=3.29, *P*>*F* 0.042, sex *P*>*F*=0.036, color *P*>*F*=0.12). Sex was only a significant factor in one model. This outcome was not an improvement over the attempt using sex alone, and no model was significant after the Bonferroni correction.

### Replication

With 89 variables there needs to be at least eight significant outcomes for the probability of more significant outcomes to drop below 0.05. The eight variables that required the fewest replicates to be significant are listed in Table 1. The maximum value in each column was the estimated sample size necessary for all eight variables to achieve significance. Only considering sex, 82 replicates are needed, while for brown psyllids 40 replicates and green psyllids would need 42. Alternatively, if significance was set at 0.000562 rather than 0.05, then the percentage phloem ingestion spent in ingestion events lasting more than 10 minutes will achieve significance first at 97 replicates per sex. If color is included in the model, then with brown psyllids one needs 26 replicates per sex to detect a difference in the percentage of phloem ingestion spent in ingestion events lasting more than ten minutes. For green psyllids, one needs 55 replicates per sex to find a significant difference in the total duration of phloem salivation when followed by phloem ingestion. Additional replication would be advised if EPG was the only data because it would be difficult to build a convincing biological narrative with a single significant outcome.

**Table 1.** The estimated number of replicates to achieve statistical significance between males and females for the eight EPG variables requiring the fewest number of replicates. Color blind assumes that color does not matter. Waveforms are xylem ingestion (G), phloem contact (D), phloem salivation (E1), and phloem sap ingestion (E2). The maximum value within each column is at the end of the table.

| Waveform | Variable Name | Color |  |  |
| --- | --- | --- | --- | --- |
|  |  | Blind | Brown | Green |
| G | Standard deviation of mean duration of G | 61 | 28 |  |
| D | Time to first sustained D from first probe | 66 | 11 |  |
|  | Number of probes after first D |  |  | 39 |
| E1 | Total duration of E1 | 82 |  | 26 |
|  | Mean duration of E1 | 73 |  | 25 |
|  | Total duration of E1 not followed by E2 |  |  | 36 |
|  | Duration of E1 event followed by first E2 lasting over 10 minutes |  |  | 42 |
|  | Duration of E1 event followed by first E2 |  |  | 31 |
|  | Total duration of E1 when followed by E2 |  |  | 23 |
|  | Number of probes after first E1 |  |  | 39 |
| E2 | Total duration of E2 | 82 | 40 |  |
|  | Mean duration of E2 |  | 33 |  |
|  | Total duration of E1 when followed by E2 plus the E2 event | 82 | 40 |  |
|  | Total duration of E1 and E2 | 59 | 35 |  |
|  | Percentage of probe duration spent in E2 |  | 30 |  |
|  | Percent E2 spent in E2 lasting over 10 minutes | 41 | 11 |  |
|  | PotE2Indx |  | 39 |  |
| Maximum |  | 82 | 40 | 42 |

### Size

We took two measures of insect size, namely head capsule width and length of metathoracic femur. Head capsule width was significantly influenced by sex (p-value=0.0086) and color (p-value=0.0144), but the interaction was not significant (p-value=0.2737) (Table 2). In contrast, the model for femur showed no significant difference for sex (p-value=0.5236), color (p-value=0.0659), or the interaction (p-value=0.2323). As expected ln(head) and ln(femur) are related (*df* 1,82, *F*=7.49 *P*>*F* = 0.0076) as head=4.3 (0.76) + 0.35 (0.13)*femur length (numbers in parentheses are standard errors), but the correlation was r^2^=0.08.

**Table 2.** Size in micrometers of psyllids based on abdominal color and sex. Means followed by different letters are significantly different by t-test. Untransformed means are shown, but analysis was done on log transformed data. We did not compare head versus femur, nor did we compare green male versus brown female or green female versus brown male. Upper case letters are differences in size of sexes while lower case are differences in size based on color.

|  |  | <u>Color=brown</u> |  |  | <u>Color=green</u> |  |  |
| --- | --- | --- | --- | --- | --- | --- | --- |
|  |  | n | Mean (SE) |  | n | Mean (SE) |  |
| Head | Female | 24 | 585.3 (4.3) | Aa | 20 | 593.3 (6.2) | Ab |
|  | Male | 26 | 593.5 (5.1) | Ba | 14 | 614.9 (7.3) | Bb |
| Femur | Female | 24 | 366.2 (2.3) | Aa | 20 | 368.1 (3.2) | Ab |
|  | Male | 26 | 363.3 (3.1) | Aa | 16 | 373.5 (1.6) | Ab |

To examine the effect of size, we started with the full model: EPG-variable=head capsule + metathoracic femur length + an interaction term. The R^2^ values in resulting models were small (<0.18) (Table 3), but there was a consistent pattern indicative of an underlying biological process. All the significant models (at the 0.05 level) have something to do with phloem ingestion. Most of them involved the time it takes to reach or begin ingesting phloem and larger individuals took longer. The exception in this trend was the time to first sustained E2 (>10 minutes) from the start of the probe with the first sustained E2. While the sign of the coefficient was negative for head capsule and was smaller than for femur length, the head capsule measure was larger than the femur and the net effect for all insects was that larger individuals take less time to reach phloem once they have found the ideal spot to ingest phloem. There was one anomaly, namely the total duration of the non-phloematic phase which was reduced with increasing size. If larger psyllids spend less time in the pathway (waveform C), xylem ingestion (waveform G), and phloem contact (waveform D) behaviors within probes and take a longer time to reach phloem ingestion one might expect that larger individuals are spending more time in non-probing because that is the only behavioral choice left. However, this was not apparent in these data. There was only one significant variable involving nutrient ingestion and it indicated that larger individuals spent more time ingesting phloem.

**Table 3.** Modeling EPG variables based on head capsule size and length of metathoracic femur showing the degrees of freedom, model F value, probability of greater F, the probability of a greater F for the independent variables and interaction term (Inter), and the coefficients for the independent variables along with the interaction term (inter_coef).

| Variable | F | P>F | R2 | Head | Femur | Inter | Head_coef | Femur_coef | inter_coef |
| --- | --- | --- | --- | --- | --- | --- | --- | --- | --- |
| DurNnprbBfrFrstE1 (2, 81) | 7.59 | 0.001 | 0.16 | 0.014 | 0.038 |  | 6.25 | 6.38 |  |
| DurNnprbBfrFrstD (2, 61) | 6.86 | 0.002 | 0.18 | 0.126 | 0.006 |  | 4.27 | 9.52 |  |
| TmFrmFrstPrbFrstD (2, 61) | 5.92 | 0.005 | 0.16 | 0.042 | 0.039 |  | 5.36 | 6.64 |  |
| TmFrmFrstPrbFrstE (2, 81) | 4.37 | 0.016 | 0.1 | 0.027 | 0.242 |  | 5.7 | 3.64 |  |
| TmFrstSusE2FrstPrb (2, 81) | 4.32 | 0.016 | 0.1 | 0.024 | 0.283 |  | 5.74 | 3.28 |  |
| TmFrstSusE2StrtPrb (2, 59) | 4.18 | 0.02 | 0.12 | 0.047 | 0.015 |  | -6.04 | 8.73 |  |
| TmFrstE2StrtEPG (2, 81) | 4.68 | 0.012 | 0.1 | 0.03 | 0.169 |  | 4.27 | 3.28 |  |
| MnDurE2 (3, 59) | 2.81 | 0.047 | 0.12 | 0.017 | 0.017 | 0.017 | 1120.67 | 1218.5 | -190.89 |
| TmFrstE2FrmFrstPrb (2, 81) | 4.16 | 0.019 | 0.09 | 0.029 | 0.27 |  | 5.41 | 3.29 |  |
| TmFrstE2FrmPrbStrt (2, 60) | 4.26 | 0.019 | 0.12 | 0.026 | 0.022 |  | -6.62 | 7.99 |  |
| TotDurNnPhlPhs (3, 60) | 2.94 | 0.04 | 0.13 | 0.024 | 0.025 | 0.024 | -361.6 | -391.9 | 61.54 |
| TmFrstSusE2 (2, 81) | 4.78 | 0.011 | 0.11 | 0.033 | 0.142 |  | 4.23 | 3.52 |  |

The acronyms are contractions of long names mostly consisting of words abbreviated as follows:
dur=duration nnprb=non-probing, bfr=before, first=first, frm=from, prb=probe, sus=sustained, strt=start, mn=mean, tot=total, nnphlphs=non-phloematic phase (waveforms C, G and D, though one could argue that D should be included)

### Published Data

The raw data and the steps for processing the data are available in the Harvard Dataverse repository. The EPG recordings were compressed before submission (https://doi.org/10.7910/DVN/FR5M53). These are processed manually to acquire quantitative data (https://doi.org/10.7910/DVN/9JLWAW). There is one data file for each insect, and these need to be combined into a single file. That file also needs to be checked for errors and modified if needed (https://doi.org/10.7910/DVN/PCIS0B). This file is used to calculate the EPG variables and to this file were added the data for sex, color, and measures of size (https://doi.org/10.7910/DVN/GYH8YW). This file is subjected to statistical analyses (https://doi.org/10.7910/DVN/ULHZSG) that are then interpreted and summarized. The SAS and R programs for doing this are included.

## Discussion

It is generally reasonable to assume that there are differences between males and females. In some cases, these differences play key roles in the ability of the vector to spread the pathogen. However, for *D. citri* the difference between the sexes in their probing behavior was small or non-existent. While we cannot prove that such differences do not exist, we did show that sample sizes would have to greatly increase to detect such the effect. An experiment with two treatments consisting of a treated group and a control group that now includes sex will need 82 replicates for each sex in both treated and control groups. If the psyllid color is included, then each treatment will have 40 of each sex for brown psyllids plus 42 of each sex for green. To put these numbers in perspective we examined 127 published manuscripts that used EPG data to get an estimate of acceptable sample sizes. The papers were all published between 1977 and 2015 inclusive. Roughly 1/3 of the published manuscripts used 10 replicates per treatment, 1/3 used 15 replicates, and 1/3 used 20 replicates per treatment. The actual number of replicates was often less than this because not all individuals perform all behaviors.

Overall, we conclude that including sex or size as an explicit part of an experimental design with this insect is not worth the additional effort necessary to reliably detect the effect. Within current sample size limitations, we expect that additional studies will occasionally find sex-associated differences but that the results will be inconsistent between studies. We suggest that the issue be revisited when dealing with pathogens because they can have a differential effect on the sexes as shown for *Bemisia tabaci* (Lu et al. 2017). While there is a risk that sex has influenced the outcome in experiments that ignore sex, experiments using only one sex describe the behavior of only half the population. Typically females are selected, but sometimes the males are more important in disease transmission (van de Wetering et al. 1998). While sex was not important there was an effect due to body size wherein larger individuals took longer to reach phloem but once there they tended to remain longer. The effect of body size was not considered in previous work looking at sex differences in this psyllid, and this effect should be considered when dealing with organisms where the sexes are of different sizes. This outcome suggests that small differences in size could have biological consequences, but the low correlation coefficients in these models suggest that the effect was minor for this insect.

## Acknowledgement

We thank Gauping Liu for technical assistance that consisted of maintaining all the plants, wiring the insects, making the recordings, and converting the recordings into data. We thank Daniel Stanton (Citrus Research and Education Center microscopy core) for assistance with the equipment and software. This research was conducted through funding received from the Citrus Research and Development Foundation.

